# The Time Scale of Evolution

**DOI:** 10.1101/464362

**Authors:** Diogo Passagem-Santos, Lilia Perfeito

## Abstract

Fitness is a measure of how quickly alleles change in frequency under natural selection. Time is always implicit in evolutionary models but its units are rarely made explicit. When measuring phenotypes such as absolute growth rate, the units of measurement need to be made explicit. By contrasting measures of fitness and growth rate, we uncovered a curious effect, by which evolutionary time runs at different speeds depending on how restricted population growth is. In other words, when the generation time of a population is externally imposed, relative fitness per generation is no longer an accurate measure of differences between genotypes. We explore this effect and describe how it affects selective sweeps, probability of fixation of beneficial mutations and adaptation dynamics. Moreover, we show that different populations cannot be compared unless they share a common reference and that our inference of epistasis can be biased by this temporal effect. Finally, we suggest less biased ways to measure selection in experimental evolution.

## I. INTRODUCTION

Selection is typically measured as the difference in fitness (when generations are discrete) or difference in log fitness (overlapping generations) (e.g. Crow and Kimura [1], Fisher [**?**]). This measure is called selection coefficient, or simply *s*. The concept of selection coefficient is extremely useful as it allows us to predict the fate of mutations under different conditions and to compare the expected contributions of natural selection and genetic drift in shaping a particular trait. The subject of selection coefficients has been extensively reviewed (e.g. [2, 3]). Of particular note is the recent work by Chevin [4] which highlights the importance of the unit in which to measure *s*.

Selection has been considered in a number of demographic ([5, 6]) and life history models ([7]). Those models describe and make predictions on the fate of mutations affecting different fitness-related traits and how they relate with the environment. However, it is always assumed that mutations affecting only growth rate behave similarly whether the population is growing or not. Any differences in dynamics are normally interpreted as changes in effective population size ([8, 9]). None of these models take into account that as the population’s mean fitness changes, the strength of external demographic restriction may also change.

In this manuscript, we explore the fate of a mutation with a fixed selection coefficient as the population increases in absolute fitness during an adaptive walk. We use the definition of selection coefficient from the classical models: the difference in log fitness between the mutant and resident alleles *s* = *ln*(*W_m_*) − *ln*(*W_r_*). We consider only mutations that affect growth rate and cases where the realized absolute growth rate of the population is imposed externally. Examples include populations that can only reach a maximum value of individuals due to a lack of space or resources, or cases where resources are supplied at a low rate. The direct consequence of this limitation is that selection appears weaker as population mean fitness increases. This effect has important consequences, particularly in the inference of selection coefficients in experimental evolution. It leads to an apparent diminishing returns epistasis as well as frequency dependent selection.

## II. THE MODEL

### 1. Definition of time units

We consider a simple growth model with continuous, overlapping generations, analogous to the one described in Crow and Kimura [1]. The overall population is modeled by

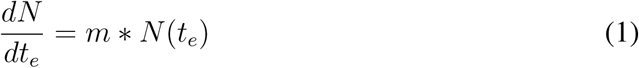

where *N*(*t_e_*) represent the number of individuals at time *t_e_* and *m* represents the Malthusian parameter of the population and is related to fitness (*W*) as *m* = ln *W*. Equation 1 represents an idealized situation that cannot exist for extended periods of time; populations have neither space nor resources to grow to infinity. Instead, they are limited and the amount of new individuals produced per unit of chronological time (*t*) in stable populations is less or equal to *m* * *N*. We interpret this discrepancy as a reduction in the effective time *t_e_* that populations spend growing. In the rest of the manuscript we will explore the far reaching consequences of this effect.

#### Continuous limitation

We consider a population that has access to a limiting resource that is constantly being renewed. At the same time a fraction of individuals die. Given a fixed rate of nutrient influx, populations are limited to produce a fixed number of individuals, *P*, per unit of *t* and die at a rate *D*. The death rate may be internally or externally imposed. Here we consider mutations that do not affect *D* as those are best described by explicit life-history models. As soon as the intrinsic growth potential of the populations surpasses the external limitation (*N*(*t*) * *m* > *P*), the population will effectively grow linearly with *t*. A practical example of such a population is the chemostat, which is a device used to continuously grow populations of microorganisms in the lab ([10]). The chemostat provides a continuous input of nutrients and a continuous removal of nutrients, by-products and cells. It is a good representation of the continuous turnover in nature [11]. Models of population dynamics in chemostat conditions usually include a concave monotonic function connecting growth rate and resource levels (see [12] for an example) and assume that equilibrium resource levels are constant and far below the half-saturation constant (see [12] and [13]). Generally, a population under such limitations can be described by

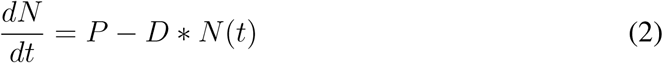

At equilibrium, 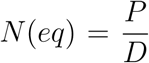. Comparing equations 1 and 2, we see that the growth component *m* * *N*(*t*) now becomes *P*. In order to relate growth in the chemostat with the classic definition of fitness, we define effective time, *t_e_*, as the time a population would spend growing in order to increase by *P*. More formally, *t_e_* and *t* can be related by comparing equation 1 with the growth component of equation 2:

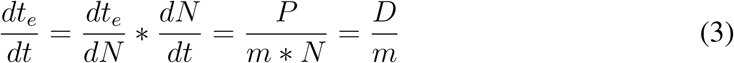

Equation 3 can be solved and a linear relationship appears between *t_e_* and 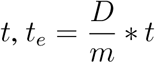. In the case of a clonal population, this is just a change of the unit in which time is measured. However, the Malthusian parameter of a population composed of different variants will be the average of the Malthusian parameters of the different sub-populations 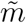. Importantly, by the action of genetic drift or natural selection, 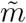 will change with time. In such populations, equation 3 becomes

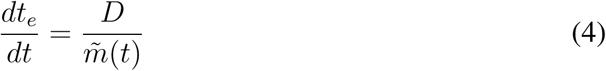

where 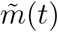 represents the mean Malthusian parameter at time *t*.

Equation 4 provides a link between chronological and effective time. The rate of effective time might not be constant and depends on *D*, which in the case of the chemostat is externally controlled.

A common normalization in evolutionary biology is to model allele frequencies using the number of generations of the population, *G*, as a time scale [12]. We define the generation time (*t_G_*) as the time it takes a population of size *N* to produce *N* new individuals, 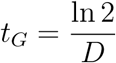 ([10]). The relationship between chronological time and number of generations (*G*) is then 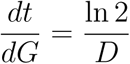. This relationship is linear as long as *D* is constant. We can now estimate how effective time and number of population generations are related,

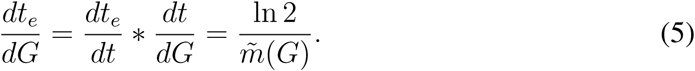

#### Discrete limitation

A different form of limitation exists when populations go through cycles of “feast and famine” whereby they receive a boost of nutrients from time to time. In between these boosts there are long periods where most of the population dies off. In this scenario, the population will grow exponentially until it exhausts all the available substrate or space and will wait for the next cycle. One example of this scenario is the widely used batch culture setting in experimental evolution ([14]). At regular intervals, these populations are subjected to a fixed bottleneck, e.g. are diluted *D′* times, and a fixed amount of substrate is made available. Between the beginning of one cycle and the beginning of the next, the population gains *P* individuals but looses a fraction *D′* of them.

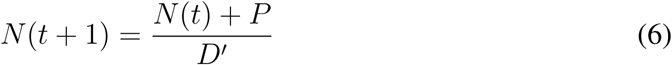

where *D′* represents the dilution factor between cycles and *P* the number of individuals produced in each cycle. Similarly to the continuous limitation, after a small number of cycles, population size at the beginning of the cycle reaches an equilibrium of *N_eq_*(0) = 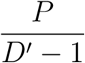. In order to estimate the effective time that passes within a cycle, we take equation 1 and substitute *N*(0) = *N_eq_*(0). We then compare the amount of growth in both models for one cycle and calculate the effective time necessary for the idealized unrestricted population of equation 1 to grow *P*:

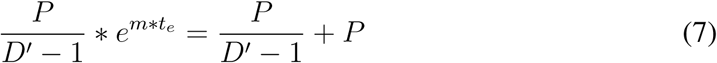

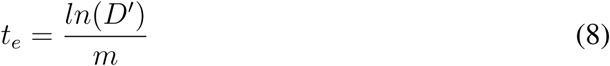

In the case of a non clonal population, equation 1 becomes:

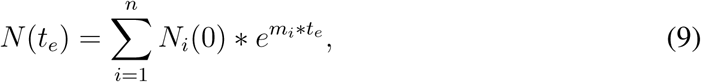

which cannot be solved analytically for *t_e_*. However, an approximation can be derived, as 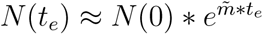, which allows us to calculate

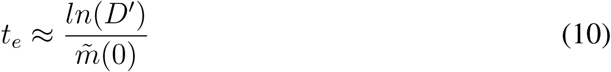

Equation 10 describes the number of units of *t_e_* in one cycle and can be generalized to *t* cycles by

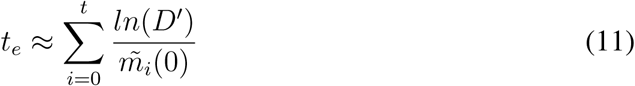

which can also be expressed in units of the number of generations of the population (*G* = log_2_ *D′* * *t*) by

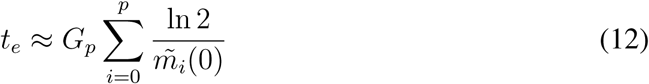

where *p* is the number of passages, *G_p_* is the number of generations per passage and 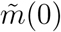 is the mean Malthusian parameter of the population at the beginning of each cycle. Comparing the results for the two types of populations limitation (equations 12 and 5) we can conclude that, when normalizing time for the number of generations of the population, the relationship between *G* and *t_e_* is the same.

#### Deterministic simulation of allele dynamics and inference of selection

In order to evaluate the analytic model in the context of competing alleles, we perform stochastic and deterministic simulations. For the case of the continuous restriction model, we make use of Monod’s equations applied to the chemostat (described in [15] and used to model evolution in [12]). For every competition we define the following system of differential equations:

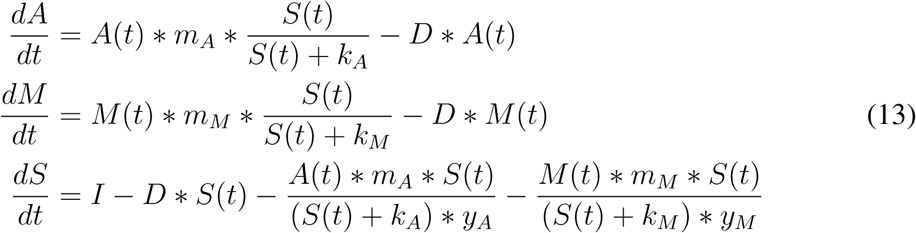

We model the ancestral strain number as *A*, the mutant as *M* and the available substrate as *S*. The dilution rate of the chemostat was defined as *D* = ln 2, with the rate of incoming substrate set as *I* = *D* * 100. Every strain is characterized with three parameters, *m* for the Malthusian parameter, *k* the half-saturation constant and *y* for the yield. We are only interested in changes in the Malthusian parameter of the populations and define *k_A_* = *k_M_* = 1 and *y_A_* = *y_M_* = 1. We are interested in simulating the dynamics of the competition starting from equilibrium conditions when the population size is constant and determined only by the substrate concentration and dilution rate. Following [16], in a chemostat population numbers 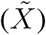 and substrate 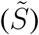 in equilibrium are given by

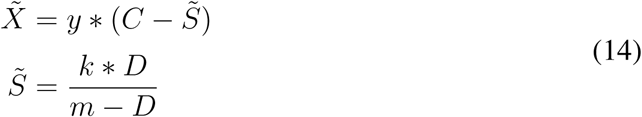

with *C* being the concentration of incoming substrate (*C* = *I/D*) and *m* being the maximum growth rate of the population. Competitions start with 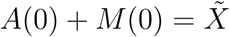 and 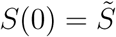.

The number of generations of the population, *G_pop_* is given by

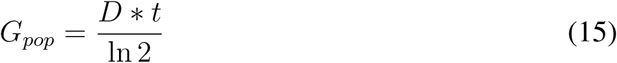

Using the system of equations 13 we estimate the allele frequencies of the population as a function of time and infer selection coefficients *s* using

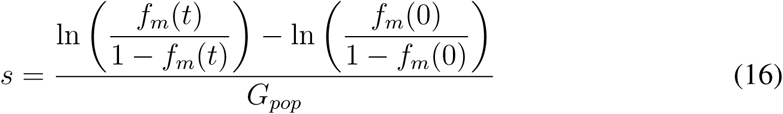

To simulate allele dynamics in a batch culture experiment (supplementary figure S1) we consider that populations can grow exponentially until the total population size is *N*(*t_e_*) = *N*(0) × *D′* and solve equation 9 for *t_e_*. The change in frequency is simply given by how much each strain grew during this time and the selection coefficient is estimated with the same equation as the chemostat. We used the deSolve package in “R: A language and environment for statistical computing” to numerically solve the system [17, 18].

#### Stochastic simulations - chemostat probability of fixation

In order to perform stochastic simulations in continuous growth restriction, we use the system of equations 13 and discretize time in intervals of about one population generation. We consider a model where at each time step one of the sub-populations is allowed to reproduce according to its growth rate and the substrate levels are adjusted accordingly (a pure jump process model [19]). A single new mutation is added once the chemostat is equilibrated 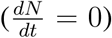 at a frequency of 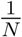. The simulations stop once one of the sub-populations is extinct. We implement it using the simulation algorithm know as Gillespie algorithm [20], in Python 2.7 [21] package StochPy [22].

#### Stochastic simulations - batch culture probability of fixation

In order to estimate the effect of growth restriction on the probability of fixation, we simulate a batch culture experiment where each sub population *k* grows deterministically following

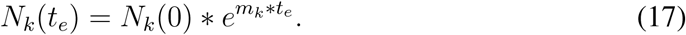

At the beginning of each passage we sample 10^6^ individuals and the population can grow until it reaches 10^8^ individuals.

Each simulation starts with a clonal ancestral population *a*, with *m_a_* = ln 2 + *s_a_*. Mutants can appear at any time during the growth cycle; to simulate the time of appearance of the mutant *t_m_* we ask in which new formed individual *X_m_* does the mutant appear by drawing from an uniform distribution between 10^6^+1 and 10^8^. Then, 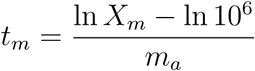. At that point we define the ancestral population as 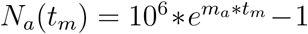 and the mutant population as *N_m_*(*t_m_*) = 1. The population can then only grow 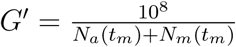 before the bottleneck. The remaining time until the bottleneck is calculated solving numerically

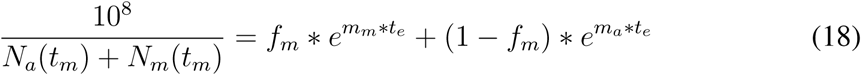

with *m_m_* = *m_a_* + *s*. The final size of each sub population is calculated using equation 17. We then sample the number of mutant individuals for the next passage *N_m_*(0) using a hypergeometric distribution and define the number of ancestral individuals as *N_a_*(0) = 10^6^ − *N_m_*(0). For every subsequent passage no new mutations are simulated. The time spent growing in each passage is calculated solving numerically equation 18. We stop the simulation once one of the sub populations disappears.

#### Simulation of a long term evolution experiment

We simulate the adaptive process in a batch culture experiment with the same general framework as described above for invasions. To introduce mutations we discretize time by a small step 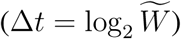 and for each strain *i* the number new mutations appearing in this time step was drawn from a Poisson distribution with average 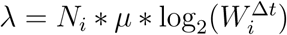, where *μ* is the per generation mutation rate. At the end of each simulated passage a new population of *N_i_* individuals is generated by sequentially sampling of each strain from a hypergeometric distribution using the final frequencies as the probability of success.

#### Simulation of discrete generations - Wright-Fisher model

In order to compare our results with classical results, we simulate populations under discrete, non-overlapping generations and constant population size. In order to compare it with the batch culture simulations, we considered an effective population size given by *N_e_* = *N_i_* × log_2_ *D′* [14]. We assume that individuals with genotype *i* produce *W_i_* × *N_i_* number of gametes per generation, such that the total number of gametes produced by the population is 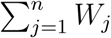. From these, *N_e_* are randomly selected to produce the next generation. The contribution of each strain for the next generation is taken from a multinomial distribution with the probability for each strain *i* being:

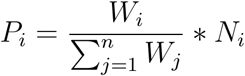

All simulations were performed in R [17] except for the probability of fixation in the chemostat, which was performed in Python, and will be made available upon request.

## III. RESULTS AND DISCUSSION

### 1. Allele Dynamics

Equations 4 and 10 show that chronological time *t* or the number of generations of the population *G* may not have a trivial relationship with the effective time the population spends growing, *t_e_*. More importantly, in non-clonal populations this relationship is non-linear and depends on the mean Malthusian parameter of the population. In this section we explore how this non-linearity can affect the dynamics of alleles. We consider a population composed of individuals carrying one of two possible alleles. We call one of these sub-populations *i* and the other the reference. We chose the reference’s doubling time as our chronological time unit, such that *m_ref_* = *ln*(2). For the moment, we assume the effect of genetic drift is negligible and consider only the changes in frequency due to selection. Following classical results ([1] and [**?**]) for unlimited populations, the frequency of sub-population *i, f_i_*(*t_e_*), is given by:

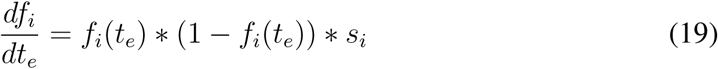

To estimate frequency changes under continuous growth limitation, we combine equations 19 and 5 to obtain the expected change in frequency of a mutant per population generation *G*:

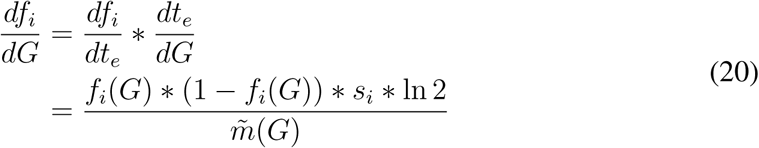

There is no closed form solution for equation 20.

In the unlimited growth model the behavior of the allele frequencies can be linearised by calculating 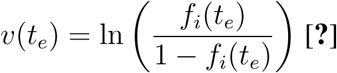. As *t_e_* and *G* do not scale linearly, *v*(*G*) should also not be linear and depend on the 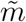 of the population. In order to test whether that is the case, we simulated a chemostat where two alleles compete under resource limitation using Monod’s growth model (see the Model section). We then compared the observed dynamics of *v*(*G*) with the predicted dynamics from the unlimited growth model (equation 19) and from the numerical solution of equation 20 (Figure 1).

**Figure 1:**
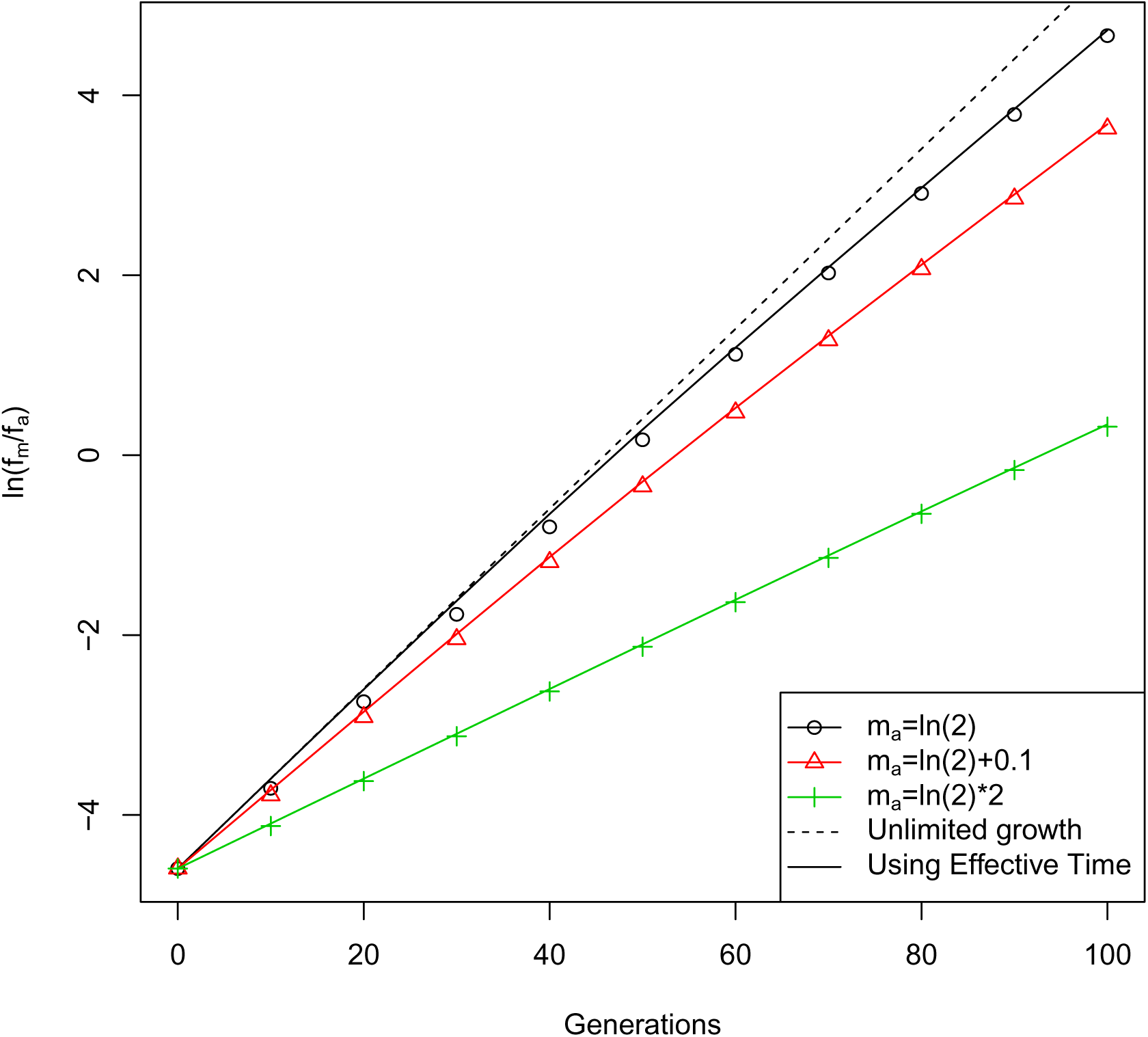
Allele dynamics with limited growth. Simulation of competition dynamics under continuous growth restriction (chemostat) with different mean Malthusian parameters but constant initial frequency (*f_m_*(0) = 0.01) and selection coefficient (*s* = 0.1). Black circles, red triangles and green crosses represent the result for the simulation of 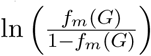 at different generations, for population with ancestral Malthusian parameters respectively of ln 2, ln 2+0.1 and ln 2*2. The dashed line represents the result of predicting dynamics using the unlimited growth model. Since s is the same for all simulations, the predicted change in frequency with this model is also the same for all three Malthusian parameters. Full lines represent the respective prediction for each simulation using the Effective Time model. *D* = ln 2.

The unlimited growth model predicts a linear relationship between *v*(*G*) and *G* [**?**], independent of the Malthusian parameter of the ancestral (dashed line in Figure 1). However, the simulated competitions display a dynamics that is not linear and depends on the mean fitness of the population (black circles, red triangles and green crosses in Figure 1). By numerically solving equation 20, we reproduce the simulated competitions (full lines in Figure 1). These results are compatible with the non-linear allele dynamics observed experimentally by [12]. The author was able to approximate the expected slope of *v*(*G*) when *f_i_* was either very small or very large. Our equation 20 can capture the whole dynamics.

The same type of analysis can be made for the case of discrete growth limitation (“feast or famine” model) and produce similar results. Solving equation 19, we get the expected mutant frequency at time *t_e_*:

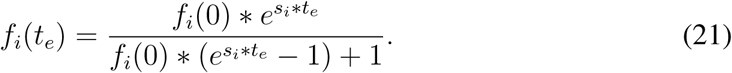

In one passage, the number of units of effective time *t_e_* that the population experiences is given by solving equation 9 for *t_e_*. For small *s_i_* we propose the approximation in equation 10. Finally, we combine equations 21 and 10 to get the expected change in mutant frequency over one passage,

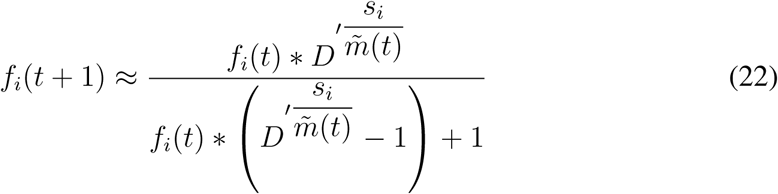

We compared the results of simulated populations under this limitation regime with the unlimited resources model and our effective time model, similarly to figure 1. We obtain the same qualitative results (supplemental figure S1). Note that the number of generations of the populations per passage is constant for constant *D′*.

Figures 1 and S1 show that under growth limitation, the trajectory of alleles under natural selection depends on the population Malthusian parameter, and not simply on the difference in Malthusian parameters of the two genotypes. However, if we are able to estimate the effective time spent growing, we can recover the results predicted by classical theory.

Another classic model in population genetics is the Wright-Fisher model [23, 24], WFM. In the WFM a population of size *N* produces an infinite pool of gametes and *N* are selected to form the new population at discrete timepoints. Given a population with two alleles, *m* and *a*, the change in frequency of *m* in one generation of the population is then defined as

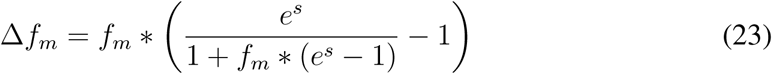

with *s* = ln *W_m_* − ln *W_a_*. Since the mean generation time in the continuous model is not constant, one generation of the WFM (*G_WFM_*) does not equal one generation in the continuous time model (*G_pop_*). In Appendix B we demonstrate that in order to observe the same change in frequency in the unlimited model and in the WFM, *G_WFM_* must equal one time unit in the unrestricted continuous model. In contrast *G_pop_* in the continuous model is the time it takes for the population to go from *N* to 2 × *N*, or 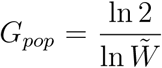. In other words,

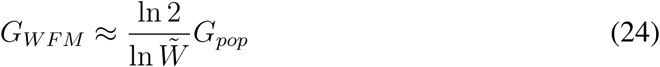

In the WFM populations express their full fitness potential in the infinite pool of gametes. This leads to a linear relationship between the number of discrete generations in the WFM and effective time in the unlimited continuous models. As seen in equation 24 the relationship between generations of a population in continuous growth and the generations in the WFM is dependent on the current mean fitness of the population.

### 2. Consequences for inference of fitness related parameters

In the previous section we show that selection coefficients based on unlimited growth do not accurately describe allele dynamics in growth-restricted populations. As a consequence, the inference of selection coefficients and other fitness related parameters will be affected. In this section we show that the change is biased, depends on the growth rate of the population and can be accounted for by measuring effective time.

#### Inference of selection coefficient

A common way to measure selection coefficients in time-series data is to use equation 19, with time measured either in absolute units (minutes, days, etc.) or in population generations. In practice, given a population composed of two genotypes, ancestral *a* and mutant *m*, the selection coefficient of the mutant (*s*) is the slope of the regression line of 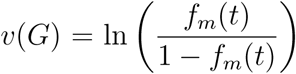 with time. As shown in figure 1, *v*(*G*) is not linear with the number of generations of the population, or with chronological time in chemostats or with the number of passages in batch culture. Therefore, the inference of the selection coefficient will be highly dependent on the initial frequency of the mutant and on the duration of the competition. In order to quantify this effect, we numerically simulated competition experiments in chemostats with different initial frequencies of mutant and different experiment duration. We then fitted a straight line to *v*(*G*) and inferred a selection coefficient. Figure 2 A shows the inference of selection coefficient for mutations with a *s* = *m_m_* − *m_a_* = 0.1. The longer the experiment is prolonged, the bigger the deviation from a straight line and the smaller the inferred selection coefficient. In the same manner, the more frequent the mutant is, the smaller is the inferred value. This apparent negative frequency dependent selection can be seen in 2B.

**Figure 2:**
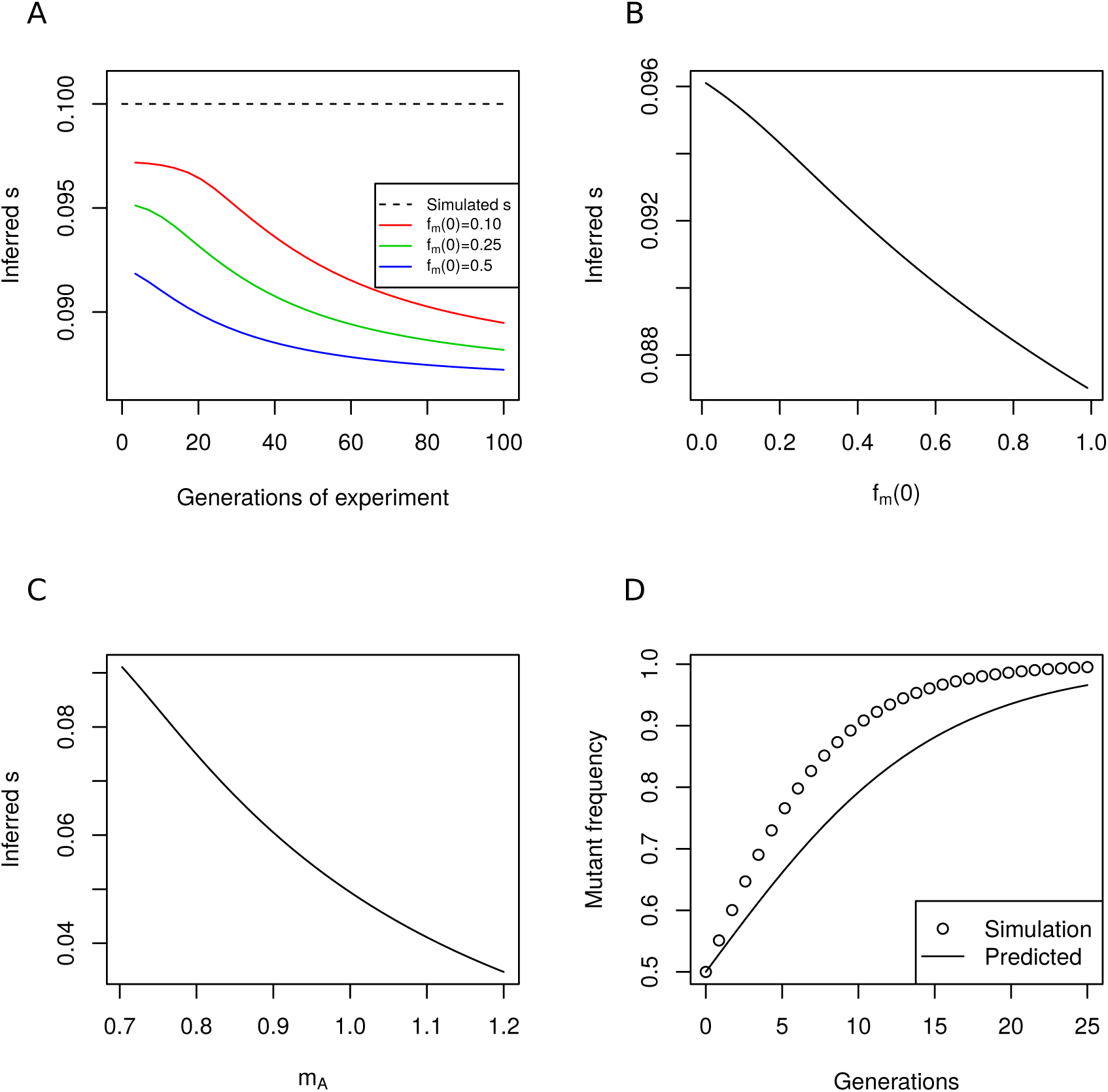
Inference of fitness related parameters. Simulation of competition dynamics under continuous growth limitation (chemostat). **A)** Inferred selection coefficient s for different duration of experiments (x-axis) and for different initial frequencies of mutant (red - 10%, green - 25%, blue - 50%). Dashed line represent the simulated s. *m_a_* = ln 2 + 0.01, *s* = 0.1. **B)** Inferred s for simulations started with different initial mutant frequencies. Simulations ran for 10 generations of the population. **C)** Inferred s for the same mutation in different backgrounds. The same reference strain was used as competitor. the selection coefficient s was estimated for different ancestral (*s_a_*) and mutant (*s_m_*) genotypes, and the s of the mutation calculated as *s_m_* − *s_a_*. **D)** Non-transitivity of selection coefficients calculated with population generations. The predicted dynamics of a competition between two genotypes, b and c (strain line) measuring their fitness versus a third genotype, a, does not follow the observed dynamics in a simulation of the competition (dots). *m_ref_* = ln 2 + 0.01, *m*_*mut*1_= ln 2 + 0.5 and *m*_*mut*2_= ln 2 + 1. In all scenarios *D* = ln 2.

#### Epistasis

One of the current challenges in evolutionary biology is to describe and understand epistasis. Epistasis exists when the effect of a mutation is dependent on the background where it appears. We define a non-epistatic system as one where the selection coefficient of a mutation is the same regardless of the ancestral genotype, such that

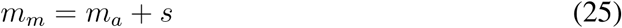

where *m_i_* is the Malthusian parameter of genotype *i*. We numerically simulate competitions with different *m_a_* and *m_m_* but the same *s*. This mimics a situation where the same mutation is put on multiple different backgrounds [**?**, 25]. All competitions were performed against a common reference (*m_ref_* = ln 2 + 0.01) and a straight line was fitted to *v*(*G*) in order to estimate *s_m_* and *s_a_*. The selection coefficient of the mutation was estimated as *s_m_* − *s_a_*. Figure 2C shows the results. For the same selection coefficient (defined in the unlimited growth model of equation 19), the more fit the background the smaller the inferred selection coefficient, leading to an apparent diminishing returns epistasis.

#### Non-transitivity

A common assumption made for mutations affecting growth rate is that they are transitive. Given three genotypes, *ref, mut*1 and *mut*2, and the selection coefficients of both *mut*1 and *mut*2 measured against *ref* (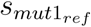 and 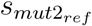 respectively), the expected advantage of *mut*2 over *mut*1 can be calculated as:

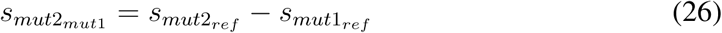

By using 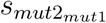 in equation 21 with *T* = *G* we can predict the dynamics of a competition between *mut*1 and *mut*2. Figure 2D shows the discrepancy between the predicted dynamics and the observed dynamics when simulating a population under growth limitation with the actual Malthusian parameters of each genotype. This is the result of the non-linear relationship between *t_e_* and *G*. Using *G* to calculate 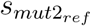 and 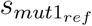 underestimates both selection coefficients in such a way that their difference is also underestimated.

#### Correction for effective time

The effects observed in the previous sections are due to the use the generation time of an evolving population as a time unit. One way around this problem is to use the generation time of one of the competitors which should be constant during the competition unless there are interactions between the genotypes. If we define the unit of time as the time the reference genotype takes to double, we can calculate the number of *t_e_* units in the experiment using only the change in frequency of the reference genotype, *f_ref_* (*t*). The reference genotype will act as our clock for measuring effective time. We start by defining *m_ref_* = ln 2 (time measured as generations of *ref*) such that,

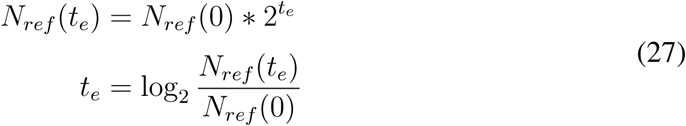

The number of reference individuals can be given by *N_ref_* (*t*) = *f_ref_* (*t*) * *N*(*t*) with *N*(*t*) representing the total population size at time *t*. The total number of generations of the population is 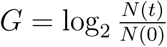, allowing us to estimate *t_e_*:

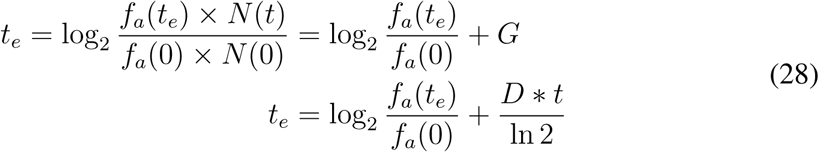

here 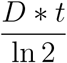 is the number of generations of the population, calculated for the continuous limitation scenario. For the discrete limitation scenario *G* = *t* * log_2_ *D′* (equation 15). The inference of *s* using *t_e_* as time unit recovers the selection coefficient from the unlimited model. With this correction, *s* is independent of both initial mutant frequency and length of the competition. Furthermore, the effect of a mutation becomes independent of the background in which it appears. We note that this is similar to the selection coefficient used by Lenski and colleagues 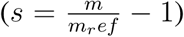 and identical to that proposed by Chevin 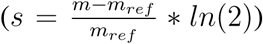. However, the use of these measurements is not widespread (e.g. [**?**]). An important consequence of using a reference as a clock for effective time is that populations that do not have any genotype in common can only be compared if we have information on the Malthusian parameters of at least one genotype per competition.

### 3. Long term dynamics

In the previous sections we describe how the inference of evolutionary related parameters is affected by the non-linear relationship between effective time and generations of the population. However, this effect is not restricted to a bias in inference. The long term evolutionary dynamics will also be affected, giving rise to non-trivial results. We will now analyse the probability of fixation of single mutations and the rate of fitness increase in populations under restricted growth.

#### Probability of fixation

We start by investigating whether growth restriction changes the probability that a mutation escapes genetic drift. This is the same as the probability of fixation in populations where the mutation rate is low enough that mutations fix one at a time. The probability of fixation of a single mutation in an haploid population (*p_fix_*) was subjected to several studies, the most notorious from Kimura (for a review on the subject see [26]). Using diffusion theory, the probability of fixation can be estimated as [27]:

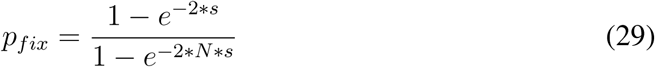

where *s* is the selection coefficient and *N* the population size. One of the assumptions of this model is that time is constant and independent of the population parameters. As we have shown before, the effective time spent competing decreases as mean population fitness increases. Given this, a mutation that appears in a fitter population (higher *m_a_*) will experience less effective time compared to a mutation with the same *s* that occurs in a less adapted population, thus having a smaller increase in frequency. The consequence is that the probability of fixation of a given mutation is dependent not only on the *s* and *N* but also on the *m_a_*.

In the case of discrete growth restriction (batch culture), equation 30 cannot be applied directly because *N* is not constant and populations go through severe bottlenecks that increase genetic drift. Wahl and Gerrish [28] studied the impact of the consecutive bottlenecks in the dynamics of new mutations and developed an equation for the probability of fixation. We follow the same methodology (more details in Appendix A) but take into account the reduction in effective time. We find that the probability of fixation becomes

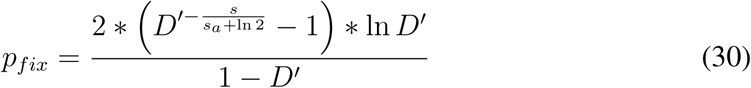

To test whether this result captures the behaviour of growth-restricted populations, we use stochastic simulations (details in the Model section above). Succinctly, a single mutation occurs at a random division in the first growth phase and no subsequent mutations are allowed. We simulate growth as a deterministic process where populations grow exponential until *N* * *D′* is achieved. We then randomly sample *N* individuals for the next passage and repeat this process until one of the strains becomes extinct. We count how often the mutant takes over the population and that is our probability of fixation. We simulate three different values of *s* for a wide range of ancestral fitness *s_a_* (Figure 3). As expected for the rationale described above, the probability of fixation decreases as the fitness of the resident population increases, and the analytic approximation presented in 30 accurately approximates the observed behaviour.

**Figure 3:**
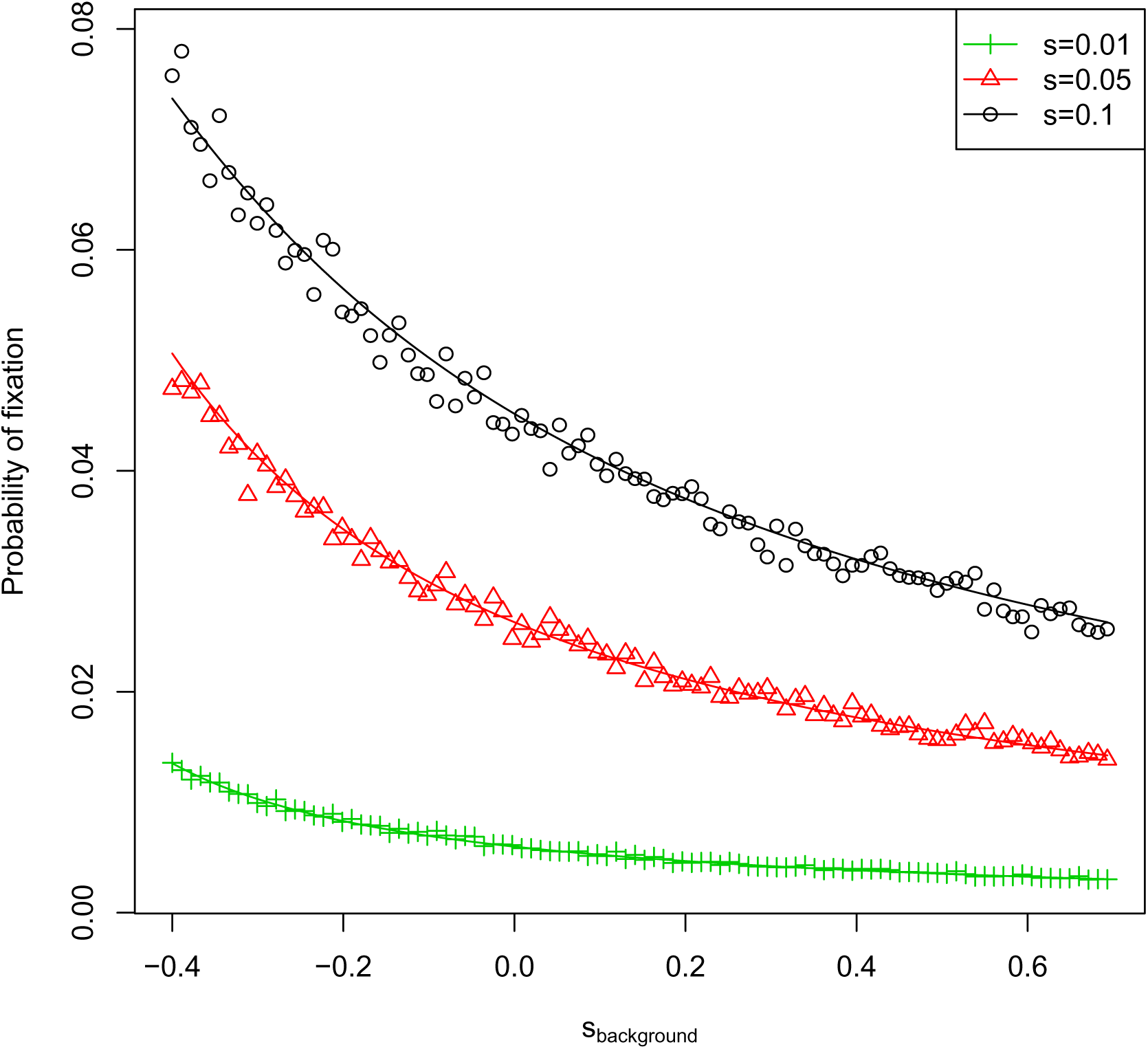
Probability of fixation in discrete limited growth. Simulated populations grow from *N_i_* =10^6^ to *N_f_* = 10^8^ individuals every passage and are randomly sampled for the next passage. The resident population has a growth rate of *m_a_* = ln 2 + *s_background_* and the mutant growth rate is *m_m_* = *m_a_* + *s*. We simulated mutations with 3 different *s*, represented in different colors. The points represent the percentage of mutations that get fixed and the lines represent the result of an analytic approximation.

The probability of fixation in populations with continuous, overlapping generations has been studied in the context of the Moran model. Briefly, in this model, the population size is constant and at every time step one individual is randomly chosen to die and another is chosen to reproduce. The probability of reproducing at each time step is given by each individual’s fitness. It has been shown that the probability of fixation in this model can be approximated by (adapted from [29]):

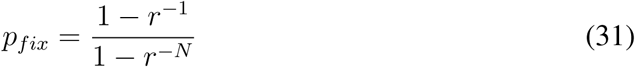

This result is obtained by defining the rates of reproduction of the ancestral *r_a_* = 1 and the mutant *r_m_* = *r*. The growth rate of each genotype is then always relative to the ancestral. Using our definitions 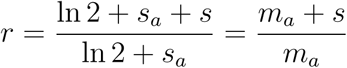. In other words, the growth rate *r* is already a relative Malthusian parameter which takes into account the ancestral growth rate. This equation accurately predicts the simulated probability of fixation in a chemostat (figure S2).

#### Rate of adaptation

In the previous sections we showed that mutations that lead to the same increase in Malthusian parameter take longer to increase in frequency as the mean Malthusian parameter of the population increases. It follows that, as populations adapt, mutations will take longer to sweep and the adaptation rate itself will slow down, even in the absence of epistasis. The rate of adaptation 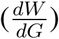 is dependent on several parameters such as mutation rate, population size and mean fitness effect of new mutations, 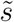. If 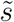 is constant, ln *W* is expected to increase linearly with time (dashed line in Figure 4).

**Figure 4:**
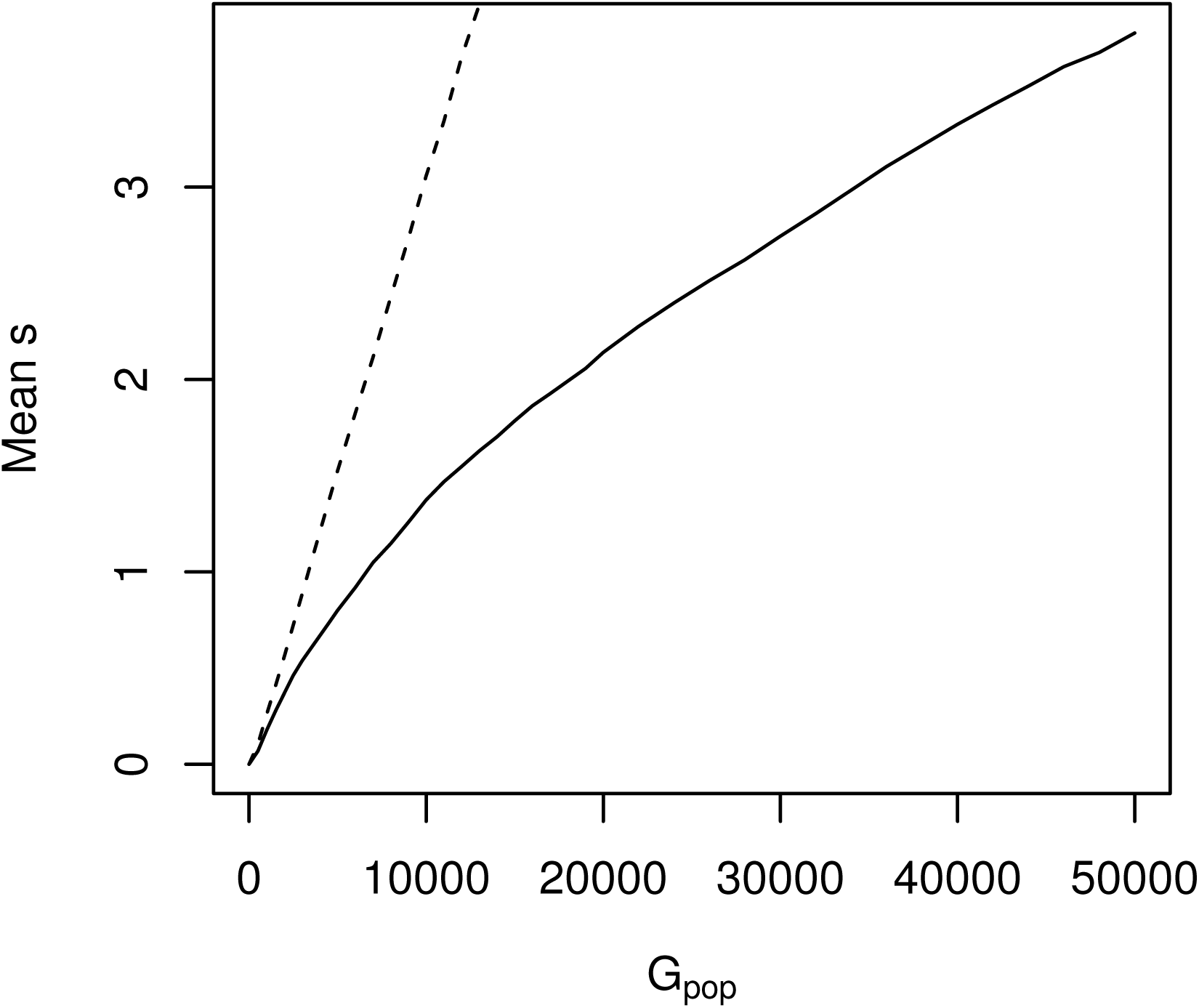
Mean log fitness in adaptation experiment. Populations were simulated with the same demographic parameters as the LTEE, *μ* = 1.7 * 10^−6^ and the selection coefficient of new mutations taken from an exponential distribution with rate λ = 100. The dashed line shows the behavior of a population of constant size and non-overlapping generations. The full line represents the simulation of a growing population with fixed *G_pop_* between passages. Each line is the mean of 12 replicates. Mean s represents 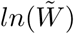 measured against the ancestral.

Given the change in time scale we report here, we expect that the rate of adaptation of the population will not be constant, but decrease as the population increases in fitness. We performed simulations of populations under growth restriction with non-epistatic mutations (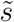 of new mutations is constant through the simulation). Succinctly, we used the same framework described above for fixation events: exponential and deterministic growth of the population until it reaches *N* * *D′* and random sampling of *N* individuals to seed next passage. In order to introduce new mutations, we discretize time in small intervals. Mutations happen at a rate which is constant per cell division and hence it is independent of *m*. We observe a pattern of decreasing rate of adaptation with number of generations of the population (Figure 4, full line). This pattern is qualitatively consistent with the observations in laboratory experiments with the same demographic parameters, like Lenski’s Long Term Evolutionary Experiment (LTEE) [14]. However, in the LTEE the rate of adaptation decelerates at a faster pace than predicted by our non-epistatic simulations. This suggests that an epistatic model is necessary to explain the observed rate of adaptation. However, not all of the decrease is due to epistasis. Next, we quantify the contribution of restricted growth in LTEE to the observed deceleration.

We start by estimating the effective time elapsed between two passages. For every two consecutive fitness measurements, *P*_1_ and *P*_2_, we approximate population growth rate by the mean of 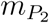 and 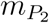. We then use equation 11 with *D′* = 100. The amount of effective time between *P*_1_ and *P*_2_ is then:

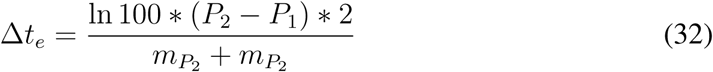

The fitness measure used in the LTEE is 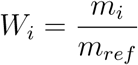. In order to obtain 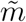, we define *m_ref_* = ln 2. By replacing *m* in the equation above, we estimate:

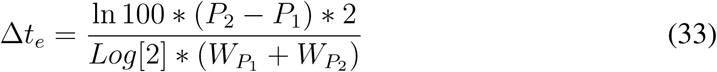

We apply equation to 33 to each of the time intervals for which there is available data. The authors in Wiser et al [30] fitted a power-law model to the data that allowed them to infer three parameters. An epistatic parameter *g*, the inverse of the mean effect of mutations in the ancestral strain *α*0 and the beneficial mutation rate *μ*. Using this approach, we fit the power law model proposed by the authors (Fig 5) to their fitness data corrected for effective time. We obtain an estimate of the epistatic parameter *g* of 5.7 (95% confidence interval 5.0 - 6.6). This estimate is within the confidence interval estimated by the authors but represents a consistent bias in the estimation of epistatic effects. The estimate of the remaining parameters was also affected, with *α*0, the inverse of the mean effect of mutations in the beginning, now 75 (95% confidence interval 51 - 146), versus the previous estimate of 85 and mutation rate of *μ* = 1.1 * 10^−6^(95% confidence interval 1.1 * 10^−6^ - 1 * 10^−5^).

**Figure 5:**
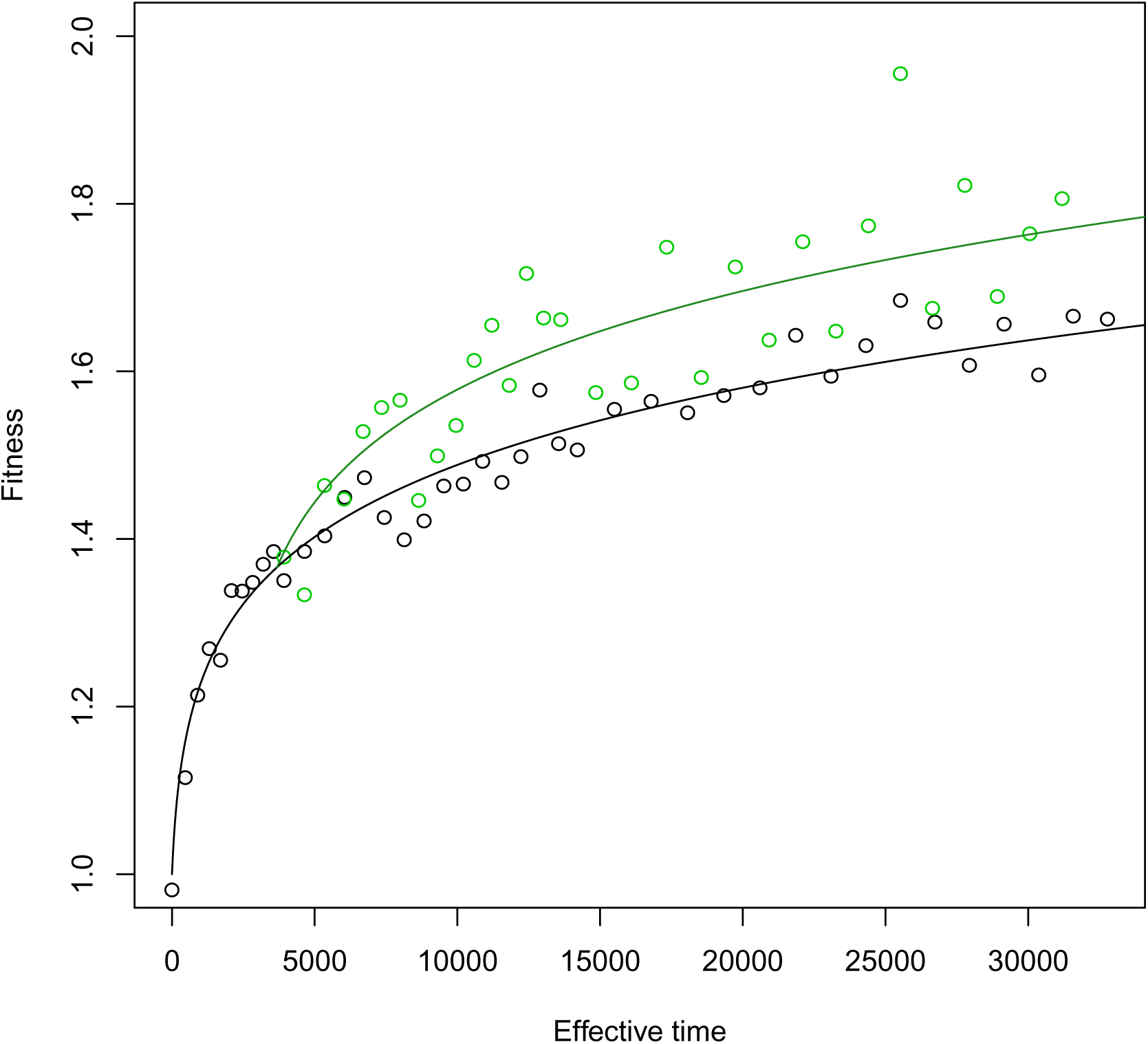
LTEE data with effective time. The black dots represent the mean fitness of the 6 populations in the LTEE that kept the initial mutation rate. Green dots represent the mean fitness of the three populations that evolved hypermutability. Effective time between two points was calculated with equation 33, after smoothing of the fitness values. The lines represent the predicted trajectory of the dynamic model with *g* = 5.7, *α*0 = 75. The black line uses a *μ* = 1.1 ∗ 10^−6^ the green line has a *μ* = 1.1 ∗ 10^−4^.

## IV. CONCLUSIONS

Fitness is often seen as an abstract term, difficult to measure for complex organisms and sometimes used tautologically. Yet, if we consider it simply as a measure of natural selection acting on a trait or genotype, fitness can be of tremendous importance in a number of contexts. For example, it can help discriminate between theories on the evolution of sexual reproduction or help us predict the probability of observing drug resistance in pathogens. In this paper, we address the strength of natural selection acting on the simplest trait: growth rate. A trait which is often equated with fitness itself. We look at populations that are growing at speeds lower than their intrinsic maximum growth rate, or Malthusian parameter *m*. This constraint probably applies to most stable populations because resources and space are finite. We show that when the population birth rate is lower than 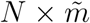, then natural selection acts more slowly that would be expected from the differences in Malthusian parameters of the different variants within the population. We interpret this reduction in speed as a reduction in the effective evolutionary time that these populations experience. We are led to the curious observation that, for the same imposed birth rate *b*, the larger the intrinsic growth rate of the population, the slower the effective evolutionary time. In other words, the faster a population is, the slower it experiences time, an affect akin to relativity in Physics.

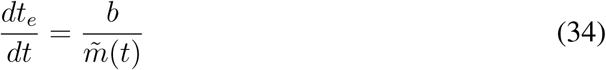

Previous work partially addressed this problem. Notably, Dean [12] showed that the selective sweep of a fitter strain in a chemostat does not follow the expected dynamics. He interpreted the discrepancy in a manner similar to ours by considering that the population mean generation time changes from the mean generation time of the resident strain to that of the fitter strain during the sweep. However, he did not fully explain the whole dynamics which we now do with equation 20. In addition he showed how this change can lead to frequency dependent selection which is able to maintain polymorphisms, which was then tested experimentally [31]. More recently, [**?**] used coalescent theory to show that the the power law behavior demonstrated in the LTEE can be attributed to a change in the time spent growing. Their approach is very different from ours but the results are consistent. However, they did not quantify the contribution of effective time to the observed LTEE results as we do. Here we go beyond specific effects of the deceleration of evolutionary time and quantify how it affects selection inference and long term adaptation.

In practical terms, this result has several implications for our understanding and expectations about the adaptation process. Mutations that are purely additive on absolute growth rate will appear epistatic on fitness measured per generation of the population. This leads to an apparent diminishing returns from beneficial mutations, a pattern often seen in experimental evolution. The reduction in effective time is not enough to explain the observations, but it likely contributes to it. As we gain more and more power to measure small selection coefficients [32], this effect will become more important. The phenomenon is not limited to measurement. If we consider that most mutations happen during reproduction, then the mutation rate will scale linearly with generation time of the population. Therefore, we expect that as a population adapts by increasing its Malthusian parameter, the time scale of mutation will become decoupled from the time scale of selection.

Here we focused on mutations affecting the growth rate only, sometimes called *r* mutations. We have not considered mutations affecting the efficiency of using resources (known as *k* mutations). There is a long literature studying the conditions under which selection is expected to be *r* or *k* driven. Among other results, it has been shown that the more constrained a population is, the stronger selection is acting on *k* mutations [33]. In our framework, this effect stems from the reduced effective time experienced by *r* mutations. We have also not investigated the behavior of mutations affecting the death rate, or frequency and density dependent mutations. The fate of all of these types will be highly dependent on the specific conditions of the experiment or population.

We have also not addressed what happens in populations with discrete generations, aside from comparing effective time with generation time in the classical Wright-Fisher model (Appendix B and equation 24). In the classical formulation of the model, at each generation, individuals are given enough opportunity to express their fitness fully because the gamete pool is infinite. Therefore, the expected relative number of offspring in the next generation for each genotype will be independent of the actual absolute fitness and environmental constraint. It will depend only on the relative fitness and frequencies of genotypes, as defined by equation 23. We have not explored what happens when the total size of the pool of gametes is externally imposed.

We show that beneficial mutations take longer to increase in frequency as their externally imposed birth rate becomes higher. This means that they spend more time at lower frequencies, where they are vulnerable to genetic drift. In turn this decreases their probability of fixation as shown in Figure 3. As a consequence, beneficial mutations behave more and more like neutral mutations as populations become more constrained. It could contribute to the amount of diversity we see in nature, which is not compatible with the selection coefficients we often measure in the lab, where populations are far less constrained.

## V. ACKNOWLEDGMENTS

We wish to thank Richard Lenski, Noah Ribeck and Mike Wiser for their helpful clarifications on the adaptation model presented in [33]. We also wish to thank the evolution community at IGC for helpful discussions. The authors acknowledge funding by the Fundação para a Ciencia e Tecnologia via PhD fellowship PD/BD/52433/2013 to Diogo Passagem-Santos, Investigador FCT IF/00019/2013 and grant EXPL/BIA-EVF/0129/2013 to Lilia Perfeito. The authors declare they do not have any competing interests.

## VI. APPENDIX A

To estimate the probability of fixation in growth restricted populations, we follow a methodology similar to the branching process described in Wahl and Gerrish [28].

Given a reference strain with *W_r_* = 2, the growth rate of the ancestral population is *m_a_* = ln 2 + *s_a_* and the growth rate of the mutant m is *m_m_* = ln 2 + *s_a_* + *s*. At the beginning of the sweep, the mean population Malthusian parameter is approximately that of the ancestral. So, the effective time in each passage is 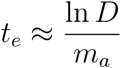, and the expected number of offspring from one mutant individual that went through a full growth phase in the next passage is 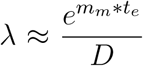. The probability *X* of this lineage to be lost in the future can be approximated using Taylor series by *X* ≈ 1 − ln *λ* This probability *X* refers to one copy at the beginning of a growth phase. The expected number of offspring of the mutant after the first bottleneck, *γ*, depends on the time of appearance *t* and is 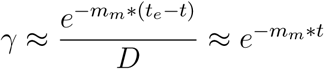. As 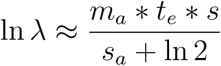, the ultimate probability of extinction (*V* (*t, s*)) is then defined as

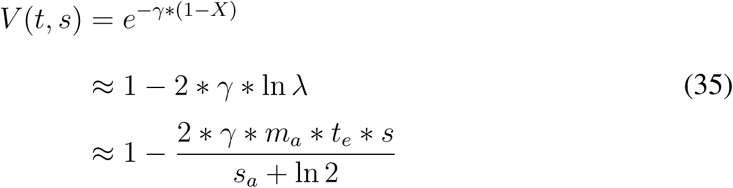

The time of appearance of the mutation is then defined as 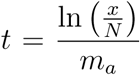, where *x* is a random variable from the uniform distribution with limits *a* = *N* + 1 and *b* = *N* * *D*. The average probability of extinction is then

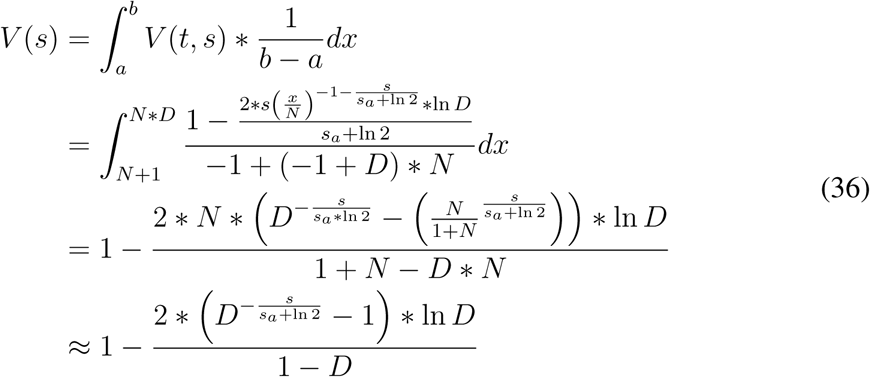

## VII. APPENDIX B

We assume a population composed of two genotypes, *a* and *m*, with absolute fitness of 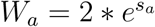 and 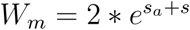. In the Wright-Fisher model, WFM, the frequency of a mutant is given by

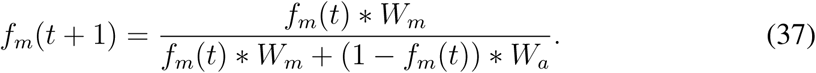

and the change in frequency in one generation of the WFM is

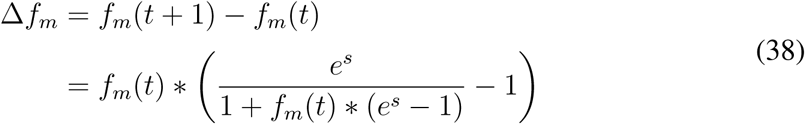

In the unlimited continuous model (equation 19) the frequency of the mutant over time can be solved as

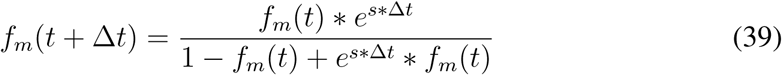

for a continuous Δ*t*. Similarly, the change in frequency of a mutant in a specific Δ*t* can be calculated as

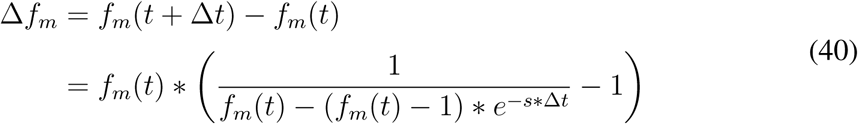

The change in frequency of the mutant is the same when equations 38 and 40 are equal

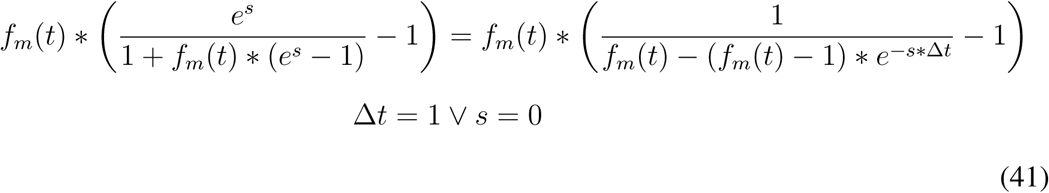

As long as the two genotypes have a fitness difference (*s* ≠ 0) the WFM and the unlimited continuous model are equivalent with a discrete step in the WFM representing a Δ*t* = 1 in the unlimited continuous model. As we show in the main text, time in the unlimited continuous model is different from chronological time in growth restricted populations. Therefore, one generation in the WFM does not equal one generation in continuous populations under growth restriction.

---

**Figure S1:**
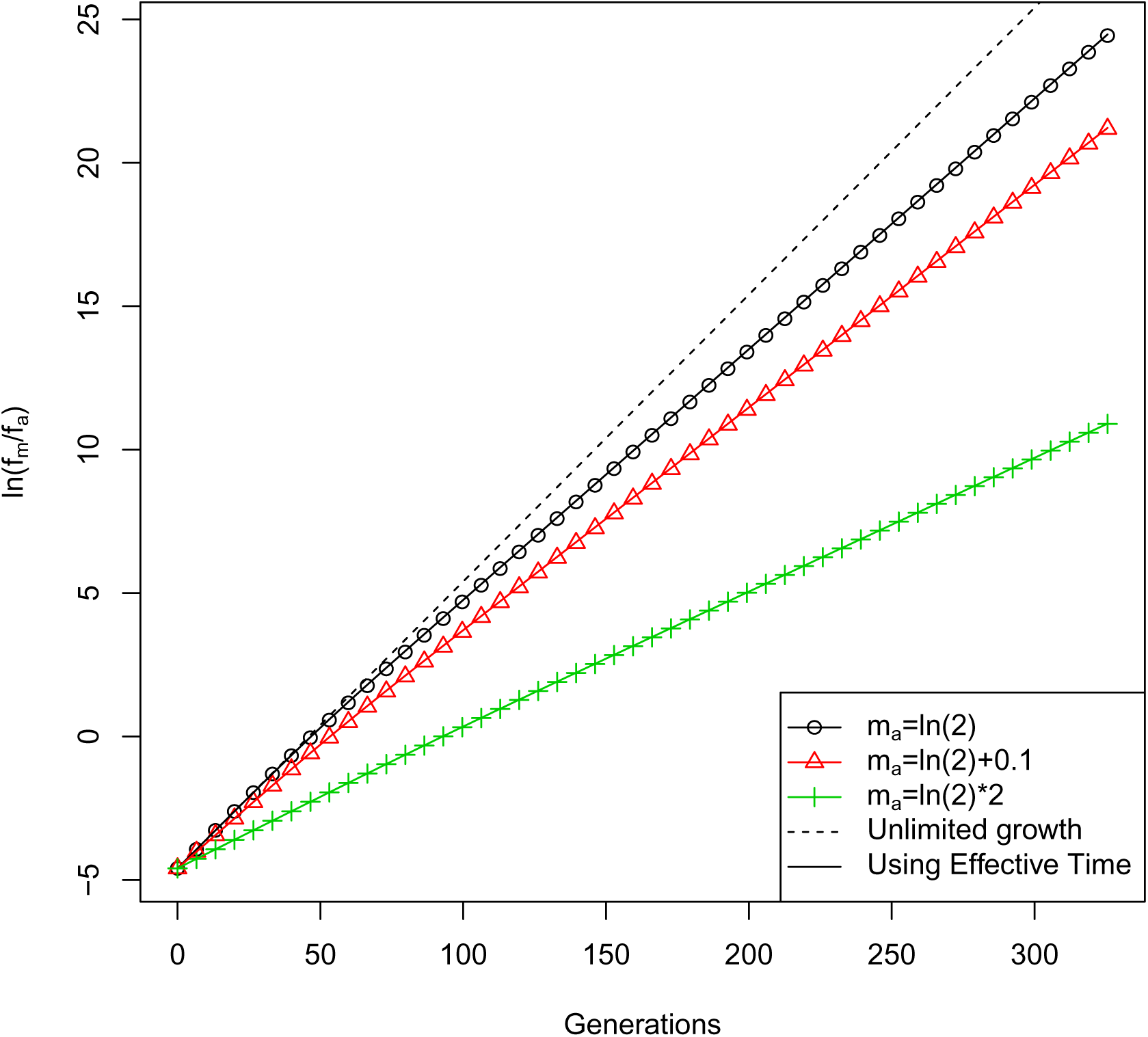
Allele dynamics with limited growth in batch culture. Simulation of competition dynamics under restricted growth with different mean Malthusian parameters but constant initial frequency (*f_m_*(0) = 0.01) and selection coefficient (*s* = 0.1). Black circles, red triangles and green crosses represent the result for the simulation of 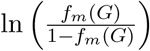 at different generations, for population with ancestral Malthusian parameters respectively of ln 2, ln 2 + 0.1 and ln 2 ∗ 2. Dashed line represent the result of predicting dynamics with unlimited growth model. Full lines represent the respective prediction for each simulation using the Effective Time model. *D′* = 100.

**Figure S2:**
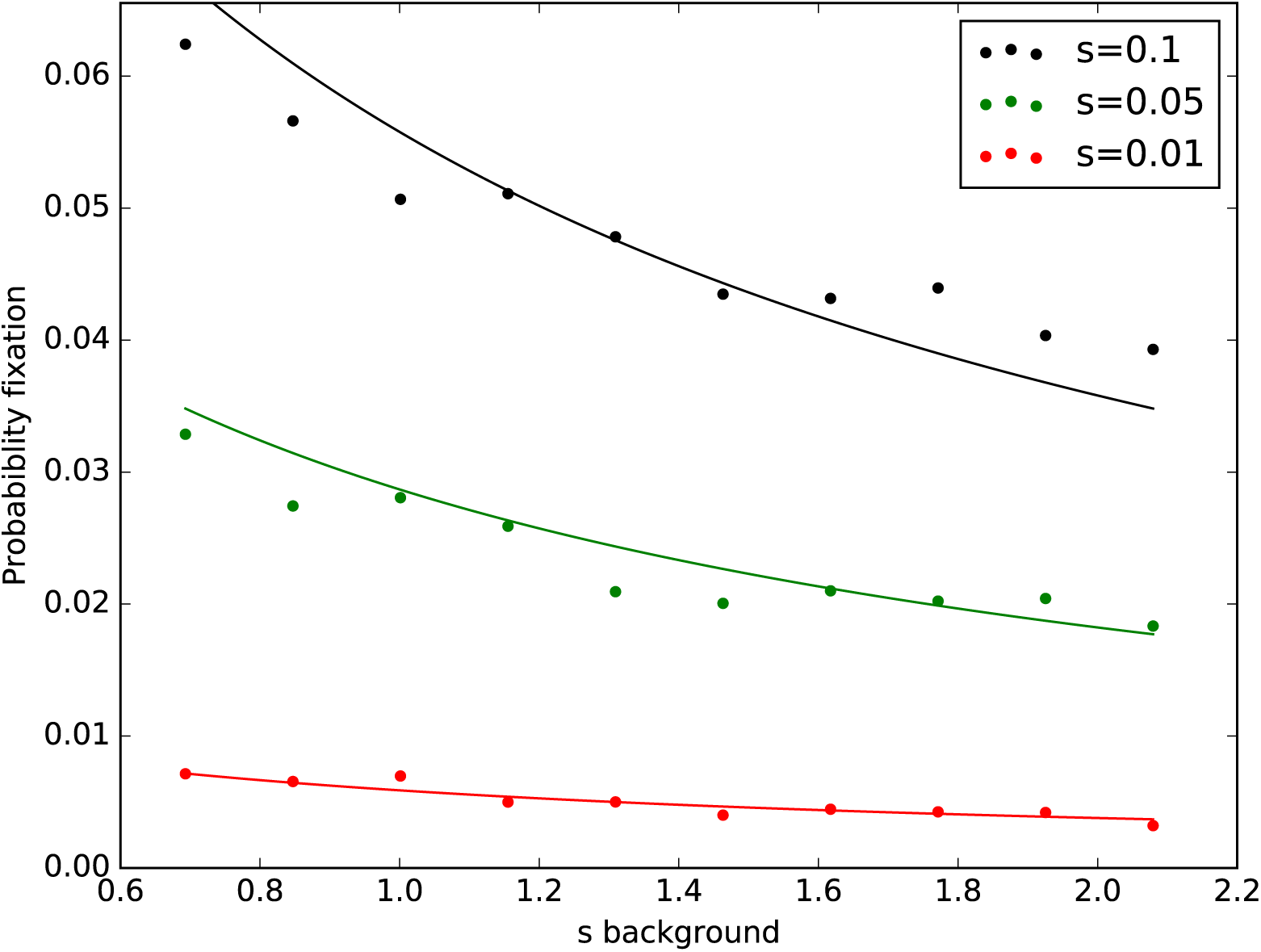
Probability of fixation in a chemostat. The dots represent the result of simulations of chemostat populations. Simulated populations start with 1000 individuals and after population reaches equilibrium a single mutant is introduced in the population. Probability of fixation is calculated as the fraction of simulations in which the mutant reaches fixation. The different background populations, represented in the x-axis, have a growth rate of ln 2 + *s_background_*. The dilution is equal in all simulations, *D* = ln 2, and the in order to keep the initial equilibrium population size constant the rate of substrate income was defined as 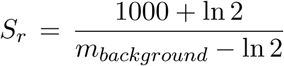. The lines represent the analytic approximation presented in equation 31 with *N* = 1000.

